# Cholinergic interneurons in the nucleus accumbens regulate the initiation of natural reward approach

**DOI:** 10.64898/2026.09.23.753195

**Authors:** Samantha O. Brown, Romain Durand-de Cuttoli, Mariam A. Mahboob, Eftychia Markopoulou, Inés Ibañez-Tallon, Jessica L. Ables

## Abstract

Despite their established role in motivated behavior, how nucleus accumbens (NAc) cholinergic interneurons (CINs) contribute to natural reward seeking remains unclear. Here, we report that NAc CINs are key modulators of the approach phase of reward pursuit. Using *in vivo* fiber photometry in *Chat-ires-Cre* mice, we show that NAc CINs are robustly activated during the approach towards several salient stimuli including food, a social target, and a familiar object. We also provide evidence that this activation is potentially driven by olfactory cues. Disrupting NAc CIN function via targeted expression of a membrane-tethered toxin results in a delayed approach towards both food and a social target, but no alterations in object investigation, general locomotion or anxiety-related measures. Together, these findings identify NAc CINs as regulators of the initiation of natural reward approach and suggest that they link sensory information with onset of motivated action.

## Introduction

The ability to seek out rewards such as food and social connection is critical for survival. Impaired motivation to engage in such reward seeking is a core feature of several psychiatric illnesses^1^. For example, major depressive disorder is commonly characterized by reduced appetite^2^ and diminished social engagement^3^. This convergence of motivational deficits across distinct reward domains suggests the existence of shared neural mechanisms that regulate the motivation to acquire different reward types. While multiple brain regions work in concert to orchestrate reward-seeking behavior, the nucleus accumbens (NAc) is a key hub, essential for integrating motivational signals and guiding approach towards rewarding stimuli^4^. In particular, the release of dopamine (DA) in the NAc plays a central role in promoting reward pursuit, as impairments in dopaminergic neurotransmission have been shown to impair approach behaviors and reward-directed learning^5^.

Cholinergic interneurons (CINs) in the NAc facilitate DA release through activation of nicotinic acetylcholine receptors on DA terminals^6,7^. While this facilitation of DA release has been consistently demonstrated *ex vivo*, how CINs contribute to reward-seeking behavior *in vivo* is not fully understood. A few studies have revealed that NAc CINs show increased activity in response to the presentation of a cue predictive of a food reward during operant tasks^8,9^. Yet, despite this clear temporal association between NAc CIN firing and reward cue presentation, manipulation of CIN activity has yielded conflicting results. Optogenetic activation of CINs in the ventral NAc shell suppresses reward seeking^10^ and decreases food consumption^8^, whereas chemogenetic inhibition has also been shown to impair reward seeking via diminished sucrose preference^11,12^ and social interaction^12^. These findings illustrate that CINs influence motivated behavior, but the precise role that NAc CINs play in the different phases of reward pursuit and across reward types remains unclear.

Here, we performed in *vivo* optical recordings of NAc CINs in awake behaving mice in a home cage environment to identify how these neurons contribute to reward approach behaviors. We then selectively silenced NAc CINs using membrane-tethered toxins targeting voltage-gated calcium channels (t-toxins)^14^ to prevent acetylcholine release and examined the effects on food consumption, social interaction and object exploration. We show that NAc CINs are activated during the approach towards rewarding stimuli such as food, a social target, and a familiar object in the homecage. We also demonstrate that silencing NAc CINs results in a delay in the approach towards food and a social target, but not a familiar object. Together, our findings establish CINs as critical regulators of the approach phase of reward-seeking behavior. These neurons likely act to fine-tune dopaminergic signaling in the NAc to drive the initiation of motivated behavior for multiple reward types.

## Results

### NAc CINs are activated during natural reward approach

To examine NAc CIN activity during reward approach, we performed fiber photometry recordings following injection of an AAV expressing jGCaMP8s in the NAc of male *Chat–ires-Cre* mice (**Figures 1A, 1B**). When mice were presented with familiar foods (chow, a banana-flavored pellet, and peanut butter cup; **Table S1**) in the homecage after an overnight fast (**Figure 1C)**, NAc CINs exhibited a significant increase in activity during the approach towards the food (**Figures 1D, 1E**). This activity peaked prior to the start of actual food consumption. The magnitude of this activation was highest for the first food reward and declined with subsequent rewards (**Figure 1L**). Similarly, when a novel, same-sex mouse was introduced into the homecage, CIN activity in the NAc robustly increased and peaked at the onset of the approach towards the social target (**Figures 1F, 1G**). These CIN responses were strongest during the first social bout and rapidly declined with repeated interaction (**Figure 1L**). Presentation of a familiar toy object elicited a comparable pattern of CIN activation during approach behavior (**Figures 1H, 1I**). Importantly, NAc CINs did not exhibit a significant change in activity during random bouts of locomotion in the absence of a rewarding stimulus (**Figures 1J, 1K**). Taken together, these observations suggest that NAc CINs are selectively engaged during the approach towards motivationally relevant targets, rather than during general movement.

**Figure 1.**
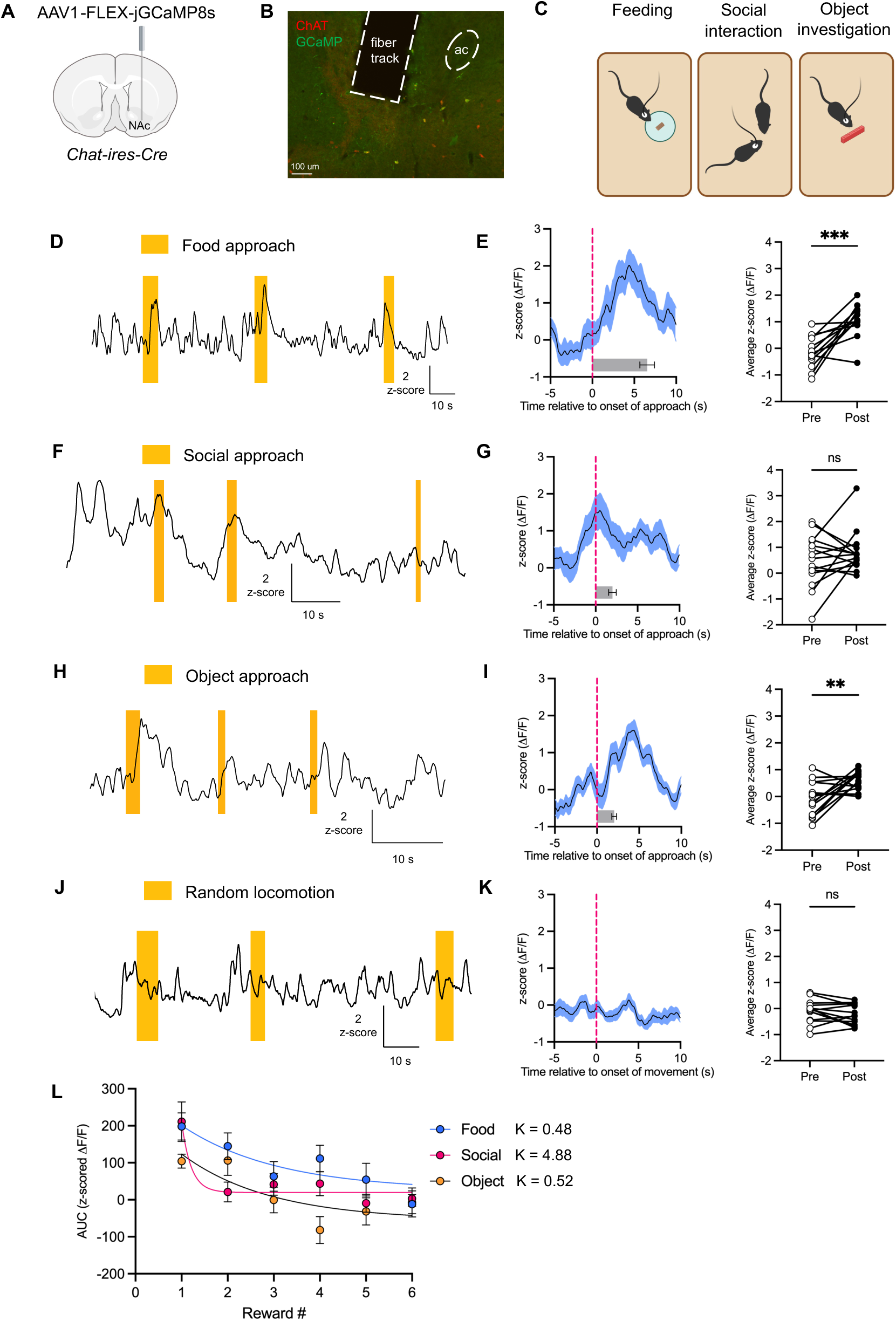
NAc CINs are activated during natural reward approach. **(A)** Schematic of unilateral AAV viral injection and fiber placement into the NAc. **(B)** Coronal brain section from *Chat-ires-Cre* mouse showing expression of GCaMP8s (stained with anti -GFP antibody) colocalized with ChAT in NAc CINs. Dashed region outlines optical fiber track. **(C)** Schematic of behavioral tests performed in the homecage. **(D)** Example trace showing NAc CIN GCaMP8s activity during feeding bouts. Yellow shaded region indicates time from approach onset to consumption onset. **(E)** Left panel shows peri-event plot of NAc CIN GCaMP8s activity aligned to onset of approach towards the first food presented (n = 13 mice). Right panel shows comparison of average z -scored ΔF/F before (-5 to 0 s) and after (0 to 10 s) the onset of food approach. Two -tailed paired t-test.^#^ **(F)** Example trace showing NAc CIN GCaMP8s activity during social interaction. Yellow shaded region indicates time from approach onset to interaction onset. **(G)** Left panel shows peri-event plot of NAc CIN GCamP8s activity aligned to onset of first approach towards a novel mouse (n = 15 mice). Right panel shows comparison of average z -scored ΔF/F before (-5 to 0 s) and after (0 to 10 s) the onset of social approach. Two -tailed paired t-test.^#^ **(H)** Example trace showing NAc CIN GCaMP8s activity during object investigation. Yellow shaded region indicates time from approach onset to object investigation onset. **(I)** Left panel shows peri-event plot of NAc CIN GCamP8s activity aligned to onset of first approach towards a familiar object (n = 16 mice). Right panel shows comparison of average z -scored ΔF/F before (-5 to 0 s) and after (0 to 10 s) the onset of object approach. Two -tailed paired t-test.^#^ **(J)** Example trace showing NAc CIN activity during spontaneous bouts of locomotion. **(K)** Left panel shows peri-event plot of NAc CIN GCamP8s activity aligned to onset of movement (n = 14 mice). Right panel shows comparison of average z -scored ΔF/F before (-5 to 0 s) and after (0 to 10 s) the onset of locomotion. Two -tailed paired t-test.^#^ **(L)** AUC of NAc CIN activity during successive bouts of reward approach in the homecage. Curves are fit to one phase exponential decay with Y _0_ > 0 constraint. K = decay constant All error bars represent ± SEM. **p < 0.01, ***p < 0.001, ns = not significant ^#^For peri-event plots, solid line indicates mean and shaded region represents SEM. Horizontal bar plot shows average time elapsed from approach onset to the onset of food intake, social contact, or object investigation.

### Olfactory detection of food and pheromone odors promotes NAc CIN activation

Given that NAc CIN activation preceded the onset of food consumption and social interaction, we hypothesized that the sensory detection of reward, through olfactory cues, might be sufficient to drive CIN activation in the absence of consummatory behavior. To test this, we exposed fasted mice to food and pheromone odors (**Figure 2A**). Consistent with our hypothesis, NAc CINs showed a significant increase in activity in response to food odors (**Figure 2C)**. Interestingly, the observed CIN activation occurred *before* the mice actually began to approach the odor applicator, suggesting a possibility that early detection of the potent food scent produced this increase in CIN activity. In addition to food odors, we also examined NAc CIN responses to a neutral water odor at the start of each recording session. While we predicted that there would be no neuronal response to a neutral odor, we observed that NAc CINs showed a robust increase in activity as mice approached the odor applicator containing this neutral odor (**Figure 2B**). This observation likely reflects that CINs are engaged in the response to novel stimuli, as the odor applicator was a novel object in the homecage for these trials.

**Figure 2.**
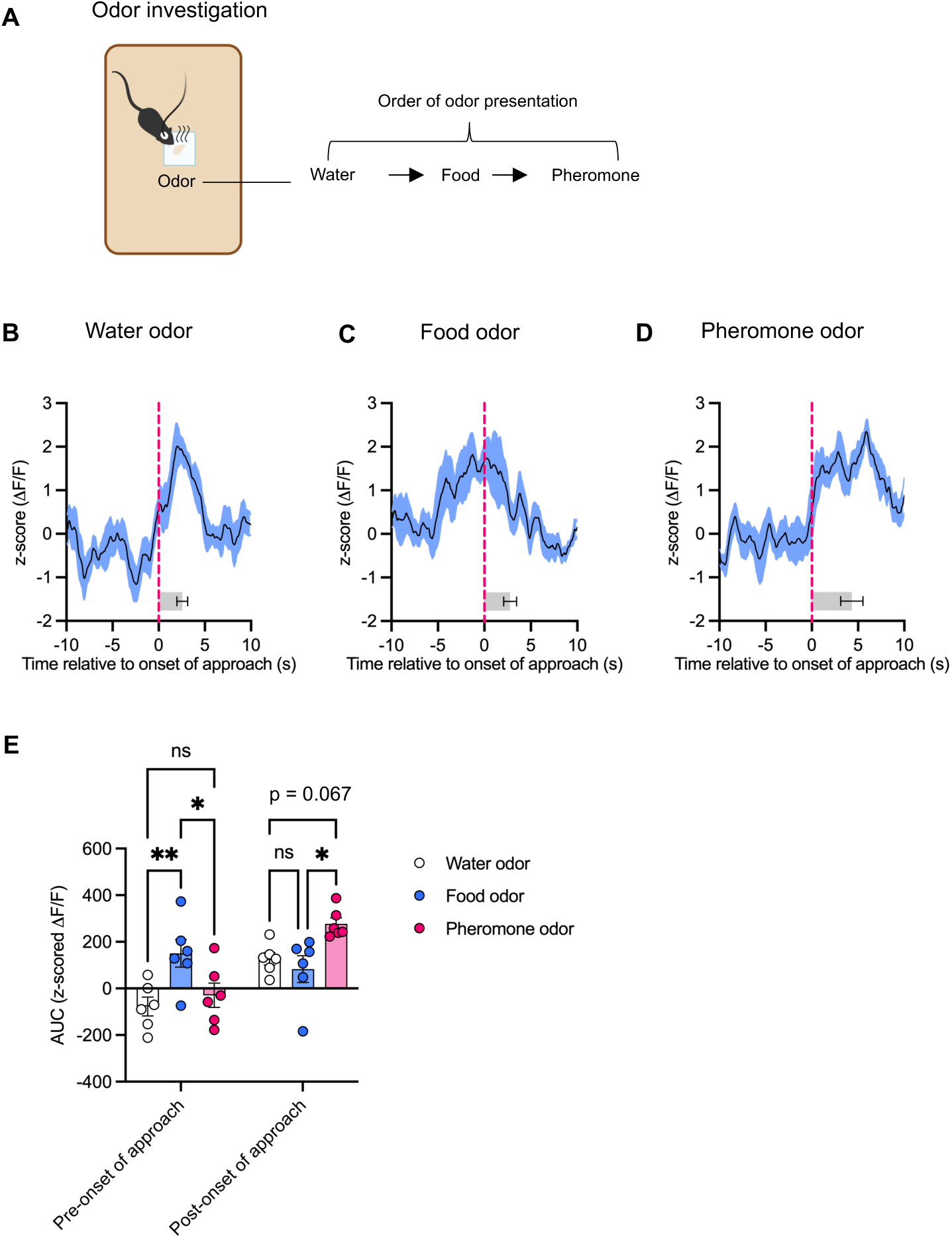
NAc CINs are activated by reward-predictive olfactory cues. **(A)** Schematic depicting odor investigation testing performed in the homecage. **(B)** Peri-event plot of NAc CIN activity during first approach towards a water (neutral) odor (n = 6 mice).^#^ **(C)** Peri-event plot of NAc CIN activity during first approach towards a food odor placed on an odor applicator in the homecage (n = 6 mice).^#^ **(D)** Peri-event plot of NAc CIN activity during first approach towards a pheromone odor (female urine) (n = 6 mice).^#^ **(E)** AUC of NAc CIN activity during 10 s before and after onset of approach towards odor applicator. Two-way ANOVA with Tukey correction for multiple comparisons. Significant main effect of time (p = 0.0027), odor (p = 0.0478), and time x odor (p = 0.006) interaction. All error bars represent ± SEM. *p < 0.05, **p < 0.01, ns = not significant ^#^For peri-event plots, solid line indicates mean and shaded region represents SEM. Horizontal bar plot shows average time elapsed from approach onset to the onset of odor contact.

To test whether NAc CINs also respond to social olfactory cues, we exposed sexually naive male mice to female urine, a pheromone odor that plays an important role in mating and is known to elicit strong investigatory behavior^13^. Remarkably, CINs in the NAc showed robust activation during the investigation of the novel pheromone odor (**Figure 2D**), even in the absence of prior mating experience. Further, during investigation of female urine, NAc CIN activity remained elevated for a longer period compared to investigation of food and water odors (**Figure 2E**). On the basis of these data, we postulate that NAc CINs are activated during approach behavior and can be activated by reward-related cues, in this case odors alone. These results highlight a potential role for CINs in integrating sensory signals to facilitate goal-directed approach behaviors.

### Silencing NAc CINs delays the initiation of natural reward approach

We next asked whether inhibition of NAc CIN activation might alter reward approach behavior. To address this question, we used a membrane-tethered toxin (t-toxin) approach^14^ to perform targeted silencing of NAc CINs (**Figures 3A, 3B**) and tested feeding behavior, social interaction, familiar object investigation, and general locomotion (**Figure 3C**). We found that NAc CIN silencing resulted in a significant delay in the approach to consume the first food reward presented in the homecage (**Figure 3D**). Interestingly, this effect was only observed for the first food reward, and not for subsequent rewards (**Figure 3E**). In addition to altered food approach, we observed that NAc CIN silencing prolonged the latency to initiate contact with a novel same-sex adult mouse in a social interaction test (**Figure 3F**), but no difference in social interaction ratio (**Figure 3G**), a general measure of sociability. Importantly, mice with silenced NAc CINs did not show increased latency to enter the SI zone during trials when the social target was absent (**Figure S1**). To investigate if the effect of NAc CIN silencing causing a delay in reward approach might be due to impaired locomotor activity, we assessed movement in an open field arena. We did not observe any effects of silencing NAc CINs on total distance travelled (**Figure 3J**) or movement velocity (**Figure 3K**) in the open field. Finally, to test if NAc CIN silencing might increase anxiety-related behavior and thus impair reward approach, we examined behavior in an open field and an elevated plus maze. We did not find any significant effect of NAc CIN silencing on the latency to enter the center of the open field (**Figure 3L**) or the open arms of an elevated plus maze (**Figure S2B**), which are both anxiogenic for rodents^15,16^. These observations demonstrates that the delayed approach behavior was not due to anxiety or to an aversion to the enclosure itself, but was rather driven by the presence of a novel mouse within the enclosure.

**Figure 3.**
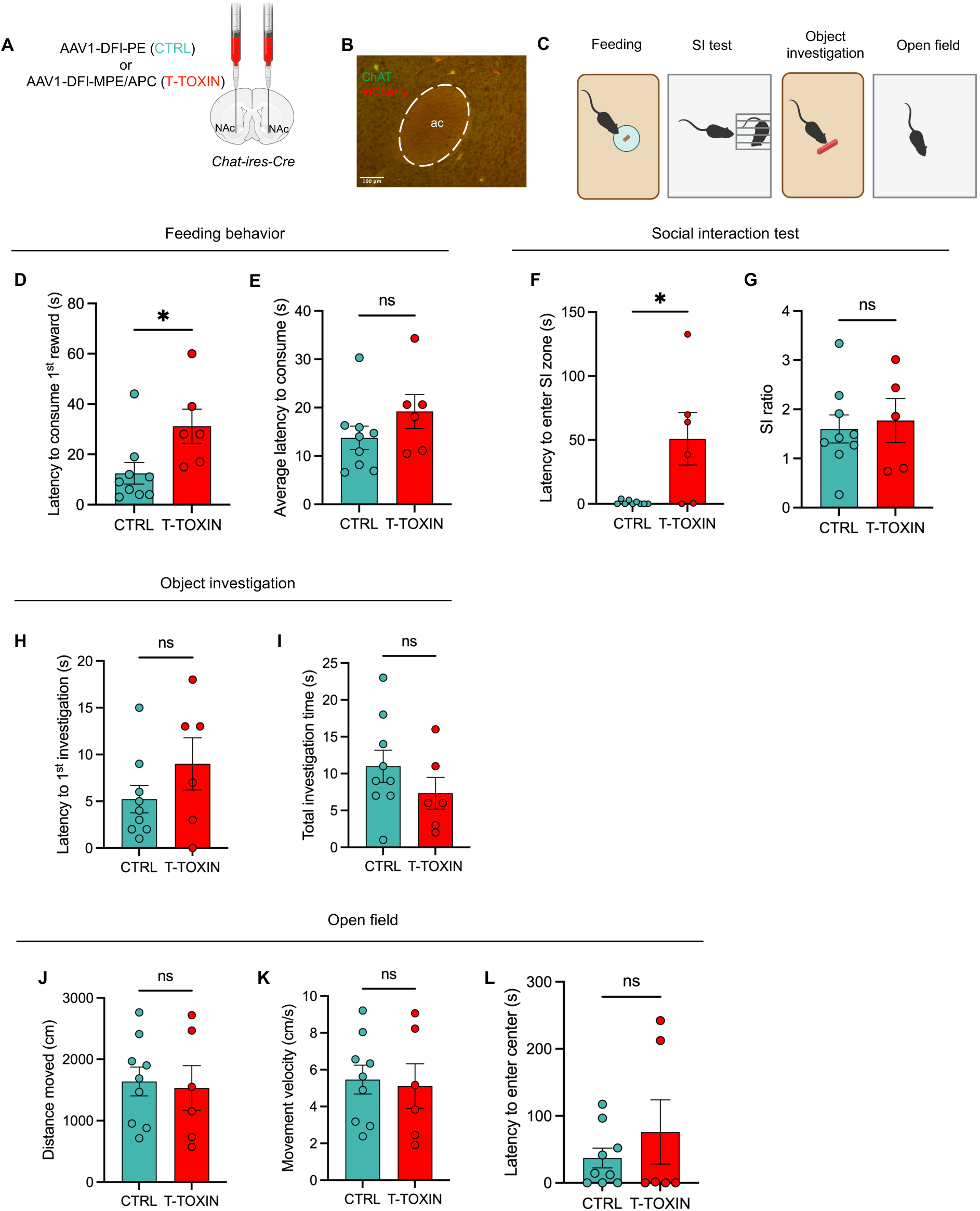
NAc CINs regulate the initiation of natural reward approach. **(A)** Schematic of bilateral AAV viral injection into the NAc. **(B)** Coronal brain sections from *Chat-ires-Cre* mice showing expression of mCherry (T -TOXIN) colocalized with ChAT in NAc CINs. No EGFP expression was detected. **(C)** Schematic of behavioral tests performed. **(D)** Latency to consume the first food reward presented in the homecage in the fasted state (n = 9 CTRL and 6 T - TOXIN). Unpaired two-tailed t-test. **(E)** Average latency to consume all food rewards presented in the homecage in the fasted state (n = 9 CTRL and 6 T-TOXIN). Unpaired two-tailed t-test. **(F)** Latency to first enter the SI zone for trials when the social target was present during the SI test (n = 9 CTRL and 6 T-TOXIN). An unpaired two-tailed Mann-Whitney U test was performed because the variances in the CTRL and T-TOXIN group were found to be significantly different using an F test. **(G)** SI ratio, calculated as the time spent in the SI zone with the target present divided by the time spent in the SI zone when the target was absent (n = 9 CTRL and 5 T-TOXIN; one t-toxin mouse (SI ratio = 33.7) was identified as an outlier using the ROUT method and was excluded). Unpaired two-tailed t-test. **(H)** Latency to first investigation of a familiar object presented in the homecage (n = 9 CTRL and 6 T-TOXIN). Unpaired two-tailed t-test. **(I)** Total time spent investigating a familiar object presented in the homecage (n = 9 CTRL and 6 T-TOXIN). Unpaired two-tailed t-test. **(J)** Total distance travelled in an open field arena (n = 9 CTRL and 6 T -TOXIN). Unpaired two-tailed t-test. **(K)** Movement velocity in an open field arena (n = 9 CTRL and 6 T -TOXIN). Unpaired two-tailed t-test. **(L)** Latency to enter the center of an open field arena (n = 9 CTRL and 6 T -TOXIN). An unpaired two-tailed Mann-Whitney U test was performed due to unequal variances in the CTRL and T -TOXIN groups. CTRL = control All error bars represent ± SEM. *p < 0.05, ns = not significant

In addition to delayed approach behavior, we also observed that NAc CIN silencing resulted in a slower rate of food consumption (**Figure 4C**) and decreased consumption overall (**Figure 4D**). Interestingly, despite the fact that CINs show a strong response during object approach using fiber photometry (**Figure 1I**), silencing these neurons did not affect the latency to initiate investigation of a familiar object (**Figure 3H**), or the total time spent investigating (**Figure 3I)**. Taken together, these data convey that NAc CINs play a crucial role in regulating the initiation of approach towards appetitive rewards (i.e. food and a novel social target) and demonstrate a critical role of NAc CINs in promoting the initiation of approach behaviors to acquire food and social rewards.

**Figure 4.**
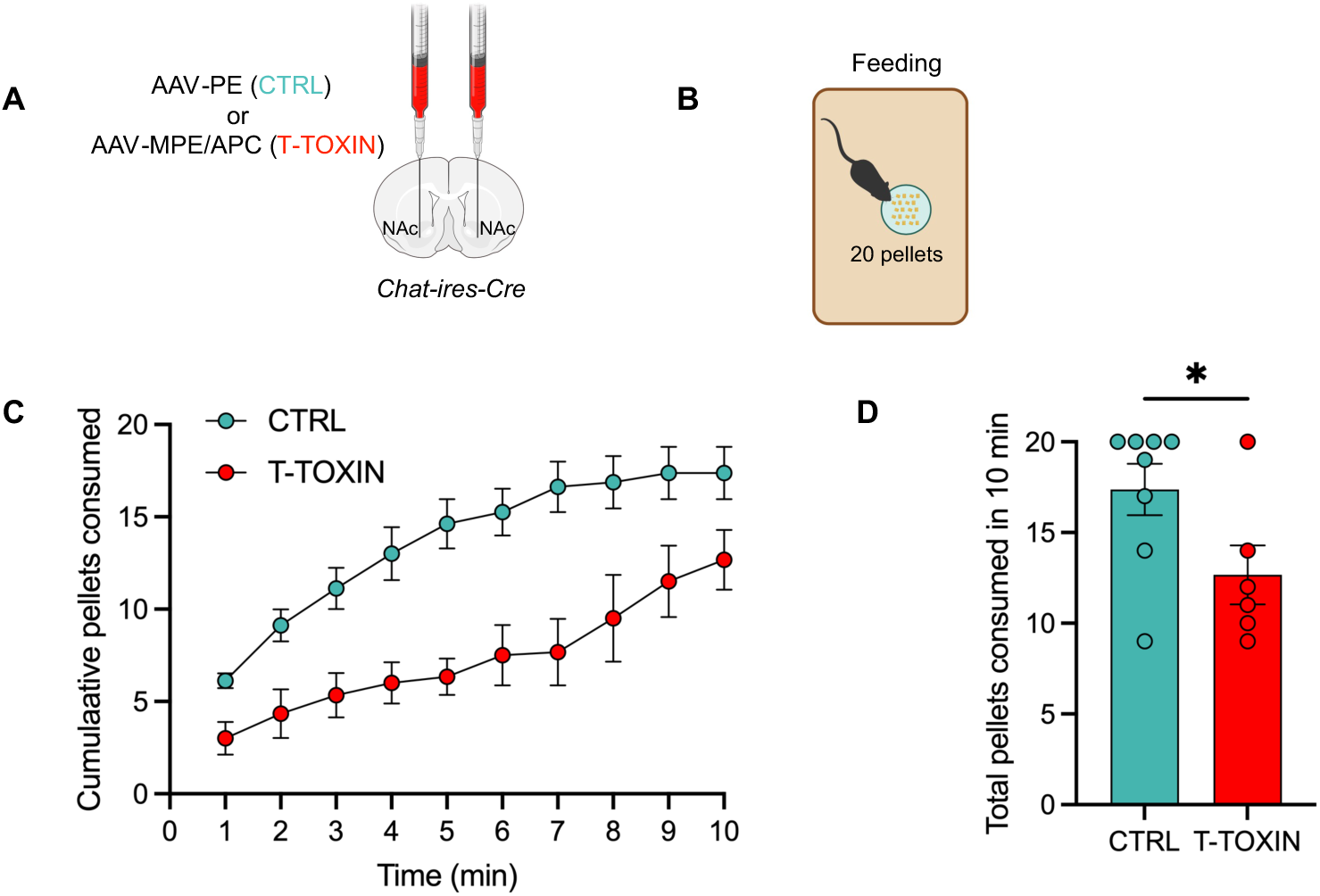
NAc CINs regulate rate and total amount of food consumption. **(A)** Schematic of bilateral AAV viral injection into the NAc. **(B)** Schematic of feeding behavior test in which mice were given 20 banana-flavored pellets in the homecage. **(C)** Cumulative number of banana-flavored pellets consumed in the homecage over 10 min (n = 8 CTRL and 6 T-TOXIN). One mouse in the CTRL group did not consume any pellets and was excluded from the analysis. Two-way ANOVA . Significant main effect of group (p = 0.0034), time (p < 0.0001), and the group x time interaction (p = 0.0006). **(D)** Total number of banana -flavored pellets consumed in 10 min (n = 8 CTRL and 6 T -TOXIN). Unpaired two-tailed t-test. CTRL = control All error bars represent ± SEM. *p < 0.05, ns = not significant

## Discussion

### NAc CINs are engaged during goal-directed reward approach

Using *in vivo* fiber photometry, we showed that NAc CINs increase activity during reward-directed approach and not during undirected movement. Notably, NAc CINs showed robust activation in response to both food and social rewards, with similar overall magnitudes but temporally distinct dynamics. CIN activity demonstrated a ramp during the approach towards food, but prior to the start of the approach towards a social target (**Figure 1**). One possible explanation is that the social target was non-stationary which may have allowed for early detection of social reward and thus an earlier rise in NAc CIN activity. Overall, our findings are in line with those of Mohebi *et al.* who showed that NAc CIN exhibit ramps in response to reward-predictive stimuli such as a cue light or the click of a food hopper^8,9^. We extend these findings by highlighting a previously undescribed role of NAc CINs during social approach behavior.

We also demonstrated that NAc CIN activation was highest for the first reward and declined across subsequent presentations of reward. This decline in CIN activation was observed for all reward types tested (**Figure 1L**), indicating that it might reflect an accommodation to repeated stimulus exposure, rather than satiety per se. Interestingly, the CIN response to a novel social target exhibited a more rapid decay compared to the responses for a familiar food or object (**Figure 1L**), suggesting that the novelty of the target might influence the rate of decline. Dopamine responses in the NAc core have also been shown to exhibit a higher decay constant for social rewards compared to objects^17^, which raises the questions as to whether CINs might be involved in modulating dopaminergic signaling in the NAc in response to novelty.

### NAc CINs are activated by olfactory detection of reward and insensitive to reward value

Our findings demonstrate that olfactory detection of reward-related odors in the environment is sufficient to elicit activation of NAc CINs. This was demonstrated using both familiar food odors and a novel social pheromone (**Figure 2**). Notably, investigation of female urine elicited significant CIN activation in sexually naive male mice. This aligns with emerging literature implicating CINs in the ventral NAc shell in the initiation of sexual behavior in male mice^18^. Furthermore, the CIN response to the novel pheromone odor remained elevated for longer compared to the response for water and food odors (**Figure 2E**), which supports the idea that these neurons might be involved in detecting the novelty of a salient stimulus, with slower decay for highly salient, highly novel stimuli.

We also investigated if NAc CINs exhibit differential responses based on the hedonic value of reward. To our knowledge, there have been no prior studies that have investigated the precise role of NAc CINs in encoding reward value. We found that the magnitude of NAc CIN activation during the approach towards a peanut butter cup, a highly preferred food (**Figure S3A**), was not higher than that for other food types (**Figure S3B**). Thus, our fiber photometry data support a model in which NAc CINs are broadly activated during the approach towards reward, regardless of type or value.

### NAc CIN activity is critical for the timely initiation of reward-seeking behavior

T-toxin viruses targeted to silence NAc CINs have previously been shown to induce a depression-like phenotype in mice characterized by reduced sucrose preference^11^. We extended these findings by illustrating a critical role of NAc CINs in modulating the timely initiation of reward approach. We show that t-toxin mediated inhibition of NAc CINs results in a significant delay to approach food and social stimuli (**Figure 3**), but does not completely abolish these behaviors, suggesting a modulatory, rather than essential, role of these neurons in approach initiation. This delayed reward approach cannot be explained by an anxiety-like phenotype, as mice with silenced NAc CINs did not show increased latency to enter anxiogenic zones on an elevated plus maze or open field (**Figures 3L, S2B**). Interestingly, mice with silenced CINs did not exhibit a significantly increased latency to investigate a familiar object compared to control mice (**Figure 3H**). This suggests that CINs are critical in regulating the approach towards rewards that enhance survival (i.e. food and social contact), although they may generally respond to other salient stimuli.

An important caveat is that while fiber photometry recordings of NAc CINs during social interaction were conducted in the homecage, the SI test for the t-toxin experiment was performed in a novel arena with the social target confined to an enclosure. This revealed a delayed social approach phenotype in mice with silenced CINs that might not have been captured if the experiment had been performed with both mice (resident and intruder) freely moving in the homecage.

On the basis of these data, we propose that NAc CINs facilitate the initiation of general reward-seeking behaviors, likely via modulation of DA release, though this warrants thorough testing. To probe this mechanism, one might inhibit cholinergic influence on DA release (i.e. through nAChR blockade on DA terminals in the NAc) and assess the effects on reward approach. Such a manipulation was performed by Mohebi *et al*. who showed that pharmacological inhibition of β2* nAChRs in the NAc increases the latency to consume a food reward^9^. While this remains to be tested for social reward, these published findings agree with our model that NAc CINs are important for the initiation of reward-seeking behaviors. Further in support of our framework, another study demonstrated that optogenetic inhibition of CINs in the ventral NAc shell increases the latency to initiate mounting, intromission and ejaculation in male mice^18^.

### Limitations of the study

This study has several limitations. First, our odor investigation paradigm does not allow precise determination of the onset of olfactory detection in mice. Future studies should employ an odor delivery system to precisely time-lock NAc CIN Ca^2+^ signals to odor sensing. Second, interpretation of the SI test in the t-toxin experiment is limited by a floor effect in the latency of the 1^st^ SI zone entry in control mice (**Figure 3F**). This arose because multiple mice were tested simultaneously, and behavior tracking with Ethovision was initiated only after all mice had been placed into the testing arenas. As a result, the first moments of social approach were not all captured. Despite this constraint, we observed a significant delay in the onset of social approach in t-toxin mice compared to controls. Finally, while we used injection coordinates to target the NAc core for the fiber photometry and the t-toxin experiments, we have yet to fully validate if our targeting was consistent across mice. Our preliminary imaging of brain sections from t-toxin mice shows that the APC, but not the MPE, construct was expressed in NAc CINs. This raises the possibility that inhibition of only one type of voltage-gated calcium channel was sufficient to produce neuronal silencing and behavioral effects.

## Acknowledgments

This study was supported by the Friedman Brain Institute at the Icahn School of Medicine at Mount Sinai and the NARSAD Young Investigator Award 28240 awarded to J.L.A. We thank members of the Ables and Russo labs for their critical reading and feedback of the experimental design and written manuscript.

## Author Contributions

J.L.A. and S.O.B. conceptualized the research questions. S.O.B. performed the stereotaxic surgeries. S.O.B. and R.D.C. conducted the fiber photometry and behavioral tests. S.O.B., M.A.M., and E.M. performed the perfusions. S.O.B. and E.M. performed the tissue sectioning. I.I.T. provided the t-toxin and control viruses. S.O.B. and R.D.C. analyzed the data. S.O.B., R.D.C., and J.L.A. contributed to writing and editing.

## Methods

### Animals

*Chat-ires-Cre* homozygous mice (B6.129S-Chat^tm1(cre)Lowl^/MwarJ; Jackson Laboratory strain #031661) were crossed with WT C57BL/6J mice (Jackson Laboratory strain #000664), and the adult (at least 8 weeks of age) male F1 offspring (*Chat-ires-Cre* heterozygous mice) were used for NAc CIN fiber photometry and t-toxin-mediated silencing. Cages of male *Chat-ires-Cre* heterozygous mice were randomly assigned to experimental groups for the t-toxin experiment. Mice were housed in a temperature- (20-22°C) and humidity- (30-40%) controlled facility on a 12h light/dark cycle (07:00-19:00) in groups of 2-5 per cage. All behavioral studies were performed during the light phase. Food (LabDiet #5053) and water were available *ad libitum* unless otherwise specified. All experimental procedures were approved by the Mount Sinai Institutional Animal Care and Use Committee and conducted in accordance with the National Institutes of Health guidelines.

### Stereotaxic surgery

Mice were deeply anesthetized with ketamine (100 mg/kg)/xylazine (10 mg/kg) and head-fixed in a stereotaxic frame (Kopf Instruments). A midline incision was made on the head to expose the skull, and a burr hole was drilled into the skull directly above the injection site. For fiber photometry experiments, a total volume of 0.3-0.4 µl of a Cre-dependent adeno-associated virus (AAV) expressing the fluorescent calcium indicator GCaMP8s (pGP-AAV1-CAG-FLEX-jGCaMP8s-WPREs; Addgene #162380-AAV1; viral titer - 1.9 x 10^13^ genome copies/mL) was unilaterally injected into the NAc using a glass micropipette (Drummond Scientific #5-000-1001-X) connected to a micromanipulator (Narishige MO-10). The virus was evenly distributed at three DV sites to ensure adequate infection of the sparse cholinergic interneurons in the NAc (AP +1 mm; ML +/- 1.3 mm; DV -4.4 -4.5 and -4.6 mm; relative to bregma). The needle was left in place for at least 5 min at each DV site. A 5 mm optical fiber with a 400 µm diameter and 0.66 numerical aperture (Doric Lenses #MFC_400/430-0.66_5mm_MF1.25_FLT) was then implanted into the NAc (AP +1 mm; ML +/- 1.3 mm; DV -4.45 mm; relative to bregma) and secured to the skull using opaque dental cement (C&B Metabond).

For t-toxin experiments, Cre-dependent AAVs expressing the membrane-tethered toxins AAV1-DFI-MPE and AAV1-DFI-APC (Rockefeller University) were combined 1:1 prior to injection. The MPE construct encodes the MVIIA conotoxin fused to the transmembrane domain of the PDGF receptor followed by EGFP.^11^ The APC construct encodes the AgaIVA agatoxin fused to the same PDGF receptor domain and mCherry.^11^ In addition, a Cre-dependent no-toxin AAV encoding the transmembrane domain of the PDGF receptor fused to EGFP was used as a control virus.^11^ A volume of 0.5 µl of virus (AAV-1-DFI-PE (control) or AAV-1-DFI-MPE/APC (t-toxin)) was bilaterally injected into the NAc and evenly distributed at three DV sites (AP +1 mm; ML +/- 1.3 mm; DV - 4.4 -4.5 and -4.6 mm; relative to bregma). The needle was left in place for at least 5 min at each DV site. To ensure adequate AAV expression, mice were allowed at least two weeks to recover before any behavioral studies.

### Food preference test

Mice were first habituated for two days to three different foods varying in nutritional content: chow (LabDiet #5053), banana-flavored full nutrition pellet (BioServ #F0079), and Reese’s snack size peanut butter cup (The Hershey Company). Mice were then fasted overnight and were acclimated to the behavioral room in dim light for at least 1 h prior to testing. During the food preference test, mice had simultaneous access to 0.5 g of each food in the homecage for 3 min. A camera was positioned above the cage to record movement and consumption events. The total amount of each food was weighed before and after the test for each mouse.

### Homecage feeding behavior

For fiber photometry studies, mice were habituated to the test foods for several days in the homecage and were fasted overnight before the experiment. On the day of recording, mice were habituated to the test room for at least 1 h in dim light. Three different foods (chow, banana-flavored pellet, and peanut butter cup) were sequentially placed into a conical tube cap at the center of the homecage. The chow and peanut butter cup were manually prepared to roughly match the size of the 20 mg banana-flavored pellet. The foods were introduced in a random order for a total of 9 rewards. Mice were allowed to consume each food, with at least 30 seconds before the next reward was introduced. If the mouse investigated a food at least twice and did not consume it, the food was removed and the next food in the sequence was placed in the cage. To reduce the likelihood that novelty might interfere with feeding behavior, the conical tube cap remained in the home cages for the entire duration of the study. Videos were captured by a camera placed directly above the homecage, and feeding behavior was manually scored. The onset of approach was defined as when the mouse elongated its body and took the first step towards the food. The onset of consumption was defined as when the mouse took the first bite of food which was determined based on head movement. For t-toxin studies, mice were habituated to the test foods for several days and were fasted overnight on the day before testing. Three different foods (chow, banana-flavored pellet and peanut butter cup) were sequentially placed into the homecage in random order, as described for the fiber photometry studies.

To examine meal consumption rate, mice were fasted overnight and were then given access to 20 banana pellets in the homecage for 10 min. Feeding behavior was captured by a camera placed above the home cage and was scored by an experimenter blind to the animal groups. The total number of pellets consumed over 10 min was recorded for each mouse.

### Homecage familiar object investigation

Mice were habituated to a plastic toy object in the homecage for several days before the test. On the day of testing, the object was placed into the center of the homecage for 2 min and movement was tracked using a video camera placed above the cage. The total time spent investigating the object (not including climbing on it) and the latency to investigate the object were manually scored by an experimenter blind to the experimental groups.

### Homecage social interaction

Social behavior in the homecage was assessed using a variation of the resident intruder test^19^. A novel *Chat-ires-Cre* adult male mouse (intruder) was placed into the homecage of the resident mouse for 2 min. Behavior was recorded using a video camera placed above the homecage. Social interaction involved sniffing, close following, and grooming of the intruder mouse. All experimental mice (both residents and intruders) were sexually naïve, and no aggressive behavior was observed.

### Homecage odor investigation

Odor applicators were constructed by taping an unscented KimWipe to a rectangular piece of cardstock (roughly 5 x 2.5 cm). Food odors were prepared by dissolving foods (chow, banana-flavored pellets and peanut butter cup) in water. Female urine was pooled from 15-20 adult female mice on the day of behavioral testing. Odors were pipetted directly onto the applicator, which was placed at the center of the homecage. Odors were presented sequentially, with water tested first as a neutral odor. Mice were allowed to investigate each odor for at least 2 min before the next odor was presented. Investigative behavior was videotaped using a video camera placed above the homecage and was manually scored by an experimenter.

### Open field test

Mice were placed into a white open-top chamber (45 x 45 x 30 cm) for 5 min. Total distance travelled, movement velocity, and time spent in the center (roughly 13 x 13 cm) of the arena were tracked using Noldus EthoVision software.

### Elevated plus maze

Mice were placed onto an elevated plus maze (arm length - 66.5 cm and height - 15 cm) for 5 min. Total distance travelled, movement velocity, time spent in the open and closed arms, and latency to enter the open arms were tracked using Noldus EthoVision software.

### Social interaction test

The social interaction (SI) test was performed as previously described^20^. Briefly, mice were placed into an open field arena containing an empty wire-mesh enclosure for 2.5 min. Mice were then removed from the arena and a novel same-sex same-strain (*Chat-ires-Cre* adult male) mouse was placed into the wire mesh enclosure. The test mouse was subsequently placed back into the arena and the total time spent in the SI zone (roughly 24 x 15 cm) around the social target as well as the latency to enter the SI zone were recorded using Noldus EthoVision software. SI ratio was calculated as the total time spent in the SI zone when the social target was present divided by the total time spent in the SI zone when the social target was absent.

### Fiber photometry and data processing

Fiber photometry was performed using a Neurophotometrics system (FP3001), according to the Neurophotometrics manual and published protocols^19,21^. One end of a fiber-optic patch cord with cubic zirconia sleeves covered with black tubing (Doric Lenses #MFP_400/430/1100-0.57_3m_FCM-MF1.25_LAF) was attached to the metal ferrule of the implanted fiber on the mouse. The other end of the cord was connected to a port on the Neurophotometrics system. Data acquisition parameters were controlled using the open-source Bonsai software (version 2.4). Fluorescence signals were simultaneously recorded from two excitation channels at 40 frames per second: 470 nm for calcium-dependent jGCaMP8s fluorescence and 415 nm as an isosbestic control to account for calcium-independent artifacts. Light at the fiber tip ranged from 40 μW to 80 μW and was constant across trials over testing days. Simultaneous recording of 40 frames per second from both the 470-nm and the 415-nm channel was achieved phase-to-phase and visualized using Bonsai. Custom MATLAB code was used to analyze the signal. To limit the influence of autofluorescence and motion artifacts, the 415 nm signal was first subtracted from the 470 nm signal. To correct for photobleaching, a double exponential curve was fitted to and then subtracted from the 470 nm signal. The relative change in fluorescence (ΔF/F) was calculated as the percentage change relative to the mean GCaMP8s signal across the entire recording session. Behavioral events, manually scored in seconds, were aligned with the fiber photometry data by multiplying the event frames by the acquisition frame rate. To perform group comparisons across mice, ΔF/F was normalized using the z-score function in MATLAB (i.e. scaling the data to have a mean of 0 and standard deviation of 1).

### Perfusion and brain tissue processing

For immunohistochemical validation of viral expression and fiber placement, mice were anesthetized with isoflurane and transcardially perfused with 30 mL of cold PBS, followed by 30 mL of cold 4% paraformaldehyde (PFA). Brains were extracted and post-fixed in 4% PFA overnight (∼18 h) at 4°C, then transferred to 20% glycerol (in PBS) for cryoprotection until sectioning. Coronal brain sections (at 40 µm thickness) were collected on a freezing microtome (Leica #SM2010 R) and stored in cryobuffer (25% glycerol and 25% ethylene glycol in PBS) at – 20°C until staining.

### Immunohistochemistry and widefield imaging

Brain sections were mounted on positively-charged glass slides and air-dried completely. To perform antigen retrieval for ChAT staining, slides were immersed in 0.01 M citrate buffer (OriGene #B05C-100B) at 95°C for 15 min, then briefly rinsed in PBS. A hydrophobic barrier (Super PAP Pen - Electron Microscopy Sciences #71310) was drawn around the sections before incubating in blocking buffer (3% normal donkey serum, 0.3% Triton X-100 in TBS) for 1 h at room temperature. Sections were then incubated in primary antibody (goat anti-ChAT 1:500 (Sigma-Aldrich #AB144P), rabbit anti-mCherry 1:2000 (Invitrogen #PA5-34974), and chicken anti-GFP 1:2000 for GCaMP and 1:500 for t-toxin and control (Aves Labs #GFP-1020) in blocking buffer overnight at room temperature. On the following day, sections were rinsed in TBS and incubated in secondary antibody (all fluorophore-conjugated antibodies were used at 1:500; Jackson ImmunoResearch) in TBS for 1 h at room temperature while protected from light. Sections were again rinsed in TBS and incubated in DAPI (0.1 µg/mL; Sigma #D9542) for 10 min at room temperature. Finally, sections were rinsed in TBS and air-dried completely prior to coverslipping with a 1.5 mm-thick glass coverslip (Fluoromount G mounting medium; Invitrogen #00495802). Slides were sealed with clear nail polish and stored at -20°C until imaging on a widefield Leica DMi8 microscope using a 10x objective.

To detect GFP in brain sections from t-toxin and control mice, signal amplification was performed. Brain sections were mounted on slides and antigen retrieval was performed in citrate buffer (OriGene #B05C-100B). Sections were then blocked for 2 h and incubated in a chicken anti-GFP antibody (1:500; Aves Labs #GFP-1020) overnight at room temperature. On the next day, sections were rinsed in TBS and then incubated in 0.3% hydrogen peroxide for 30 min at room temperature to quench endogenous peroxidases. Sections were then rinsed in TBS and incubated in a biotinylated anti-chicken secondary antibody (1:200; Jackson ImmunoResearch #703-065-155) for 1 h at room temperature. Sections were then rinsed in TBS and incubated in an avidin-biotinylated (ABC) HRP complex (Vector Laboratories #PK6100) for 1 h at room temperature. Sections were rinsed again in TBS. Finally, sections were incubated in cyanine 5 tyramide buffer (1:50 in 1X amplification diluent; APExBIO #K1052) for 10 min at room temperature before rinsing in TBS, incubating with DAPI, and coverslipping.

### Statistical analyses

All details of sample sizes and statistical analyses performed can be found in figure legends. Data were processed and analyzed using MATLAB R2023b and GraphPad Prism 10.4.1. For comparisons between two independent groups, either a two-tailed t-test or a nonparametric Mann-Whitney U test (in the case of non-normal data distributions) was performed. For comparisons involving three or more groups, a one-way or two-way ANOVA was used. A significance threshold of ɑ = 0.05 was used for all statistical tests.

### Reproducibility

Fiber photometry experiments were performed in two separate cohorts, with the exception of the odor testing being performed in only one cohort. The t-toxin experiment was conducted in one cohort.

**TABLE 1:** Key resources table.

| REAGENT or RESOURCE | SOURCE | IDENTIFIER |
| --- | --- | --- |
| **Antibodies** |  |  |
| Goat anti-ChAT | Sigma-Aldrich | Cat# AB144P |
| Chicken anti-GFP | Aves Labs | Cat# GFP-1020 |
| Rabbit anti-mCherry | Invitrogen | Cat# PA5-34974 |
| Alexa Fluor 647 donkey anti-goat | Jackson ImmunoResearch | Cat# 705-605-147 |
| Alexa Fluor 488 donkey anti-goat | Jackson ImmunoResearch | Cat# 705-545-147 |
| Alexa Fluor 488 donkey anti-chicken | Jackson ImmunoResearch | Cat# 103-545-155 |
| Cy3 donkey anti-rabbit | Jackson ImmunoResearch | Cat# 711-165-152 |
| Biotinylated donkey anti-chicken | Jackson ImmunoResearch | Cat# 703-065-155 |
| **Bacterial and virus strains** |  |  |
| pGP-AAV-CAG-FLEX-jGCaMP8s-WPRE | GENIE Project | Addgene:162380-AAV1 |
| AAV1-DFI-MPE | Rockefeller University | N/A |
| AAV1-DFI-APC | Rockefeller University | N/A |
| AAV1-DFI-PE | Rockefeller University | N/A |
| **Chemicals, peptides, and recombinant proteins** |  |  |
| DAPI | Sigma-Aldrich | Cat# D9542 |
| ABC-HRP kit | Vector Laboratories | Cat# PK6100 |
| Cy5 TSA fluorescence system kit | APExBIO | Cat# K1052 |
| Citrate buffer | OriGene | Cat# B05C-100B |
| Hydrogen peroxide | Sigma-Aldrich | Cat# H1009 |
| **Experimental models: Organisms/strains** |  |  |
| Mouse: C57BL/6J | The Jackson Laboratory | Stock #000664 |
| Mouse: B6.129S-Chat<sup>tm1(cre)Lowl</sup>/MwarJ | The Jackson Laboratory | Stock #031661 |
| **Software and algorithms** |  |  |
| Bonsai | Neurophotometrics | 2.4 |
| MATLAB | Mathworks | R2023b |
| GraphPad Prism | GraphPad Software | 10.4.1 |
| EthoVision XT | Noldus | Version 15 |
| Fiji (ImageJ) | Schindelin *et al.*<sup>9</sup> | N/A |
| **Other** |  |  |
| Optical fibers | Doric Lenses | N/A |
| Fiber photometry rig | Neurophotometrics | FP3001 |
| Widefield microscope | Leica Microsystems | DMi8 |
| Microtome | Leica Microsystems | SM2010 R |
| Banana-flavored pellets | BioServ | Cat# F0079 |
| Reese's snack size peanut butter cups | The Hershey Company | N/A |

