## Supplemental Figure 1 for "Cholinergic interneurons in the nucleus accumbens regulate the initiation of natural reward approach"

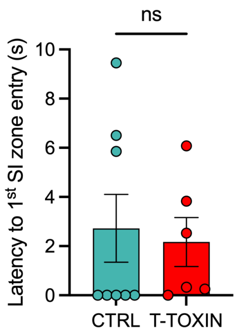


**Figure S1. Silencing NAc CINs does not affect latency to enter SI zone in the absence of a social target**

Latency to enter the SI zone during trials when there was no social target in the enclosure.

Unpaired two-tailed t-test. One CTRL mouse was identified as an outlier using the ROUT method (latency = 37.6 s) and was thus excluded from the analysis (n = 8 CTRL and 6 T-TOXIN).

CTRL = control, SI = social interaction

ns = not significant

Error bars represent ± SEM.
