## Supplemental Figure 2 for "Cholinergic interneurons in the nucleus accumbens regulate the initiation of natural reward approach"

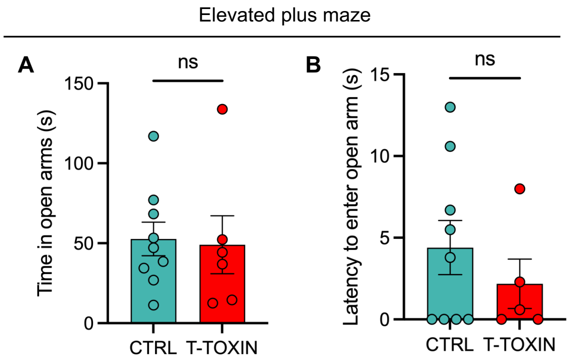


**Figure S2. Silencing NAc CINs does not alter anxiety-like behavior**

**(A)** Time spent in the open arms of an elevated plus maze (n = 9 CTRL and 6 T -TOXIN). Unpaired two-tailed t-test.

**(B)** Latency to enter the open arms of an elevated plus maze (n = 9 CTRL and 5 T -TOXIN). Unpaired two-tailed t-test. One T-TOXIN mouse was identified as an outlier using the ROUT method (latency = 92.7 s) and was excluded from the analysis.

CTRL = control

ns = not significant

Error bars represent ± SEM.
