## Supplemental Figure 3 for "Cholinergic interneurons in the nucleus accumbens regulate the initiation of natural reward approach"

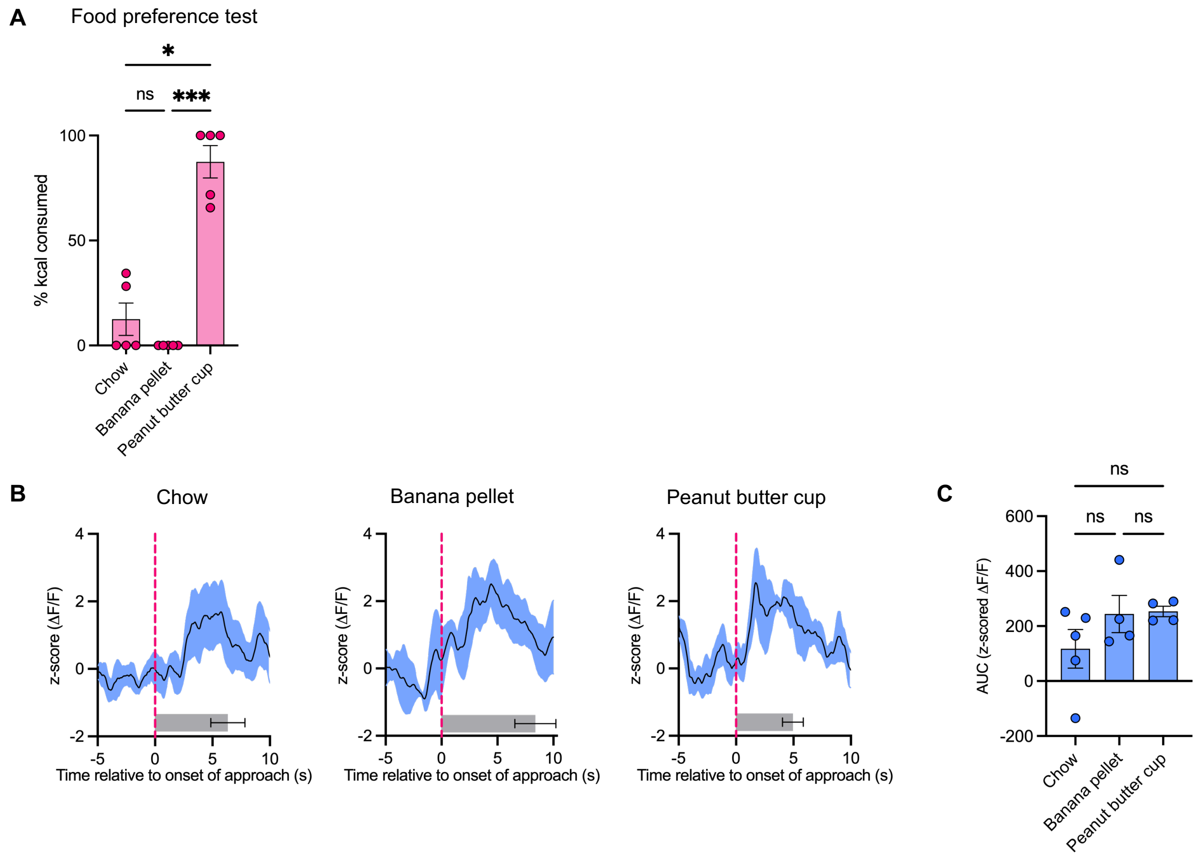


**Figure S3. NAc CIN activity is not influenced by food palatability**

**(A)** Percent calories consumed of each food type during a 3-choice food preference test in the homecage (n = 5 mice). Repeated measures one-way ANOVA with Tukey correction for multiple comparisons.

**(B)** Peri-event plots showing NAc CIN GCamP8s activity aligned to onset of approach towards first food reward in the homecage (chow n = 5, banana n = 4, and peanut butter n = 4 mice).[#]

**(C)** Area under the curve of NAc CIN activity during -5 to +10 s time window around onset of first approach towards food reward. One-way ANOVA with Tukey correction for multiple comparisons.

All error bars represent ± SEM.

*p < 0.05, ***p < 0.001, ns = not significant

[#]For peri-event plots, solid line indicates mean and shaded region represents SEM. Horizontal bar plot shows average time elapsed from approach onset to consumption onset.
