## Supplemental Table 1 for "Cholinergic interneurons in the nucleus accumbens regulate the initiation of natural reward approach"

| Food | kcal/gram | Protein % | Fat % | Carbs % |
|---|---|---|---|---|
| Chow | 4.11 | 24.5 | 13.1 | 62.4 |
| Banana pellet | 3.45 | 23.5 | 16.2 | 60.3 |
| Peanut butter cup | 5.24 | 6.25 | 56.25 | 37.5 |

**Table S1. Nutritional content of foods used in behavioral studies**
